# Ethics review for insect research in the UK: Patterns, drivers, and perceived impact on research quality and process

**DOI:** 10.1101/2025.10.21.683772

**Authors:** Jessica Eleanor Stokes, Bob Fischer, Meghan Barrett

**Affiliations:** Royal Entomological Society, The Mansion House, St Albans, UK; ProScience Ltd, Dursley, Gloucestershire, UK; Department of Philosophy, Texas State University, San Marcos, USA; Department of Biology, Indiana University Indianapolis, Indianapolis, USA

**Keywords:** Insect, ethics, welfare, insect research, 3Rs, entomology, UK

## Abstract

Research involving insects currently falls outside the scope of animal welfare legislation in the UK; however, many institutions in the UK require submission of procedures involving insects to their internal ethics review board. Here, we have surveyed UK institutions to assess what proportion require ethics applications for work with insects. Further, we surveyed entomologists working at UK institutions to understand when requirements first came about at their institution and (if applicable) their experiences with ethics review, including feedback received, resources used/desired, and their self-reported confidence in submitting applications. In addition, we assessed entomologists’ perception of the influence of ethics review on research quality and their general attitudes and values towards the ethical treatment of insects in research and the process of ethics review. Our data demonstrate that just over half of UK institutions (51.6%) require ethics review for research with insects starting in 2008 with a steady trend towards increasing institutional requirements over time. Entomologists currently submitting are largely confident in their applications (68.8%, confident) but the majority believed that resources like training on ethics in the insect context and greater research on insect welfare would be helpful. Entomologists largely believed that ethics applications had either a positive effect (44.6%) or no effect (39.6%) on research quality; early career entomologists were more likely to report a positive effect of ethics review on study quality. Qualitative responses suggest the entomological community has diverse values and attitudes towards the ethical treatment of insects in research.

## Introduction

Research involving any ‘protected’ animal species in the UK, which includes all live vertebrates and cephalapods, falls under the Animal Welfare (Scientific Procedures) Act 1986 (ASPA), a piece of legislation that regulates procedures that may cause suffering, pain, distress or lasting harm. Research involving insects currently falls outside the scope of ASPA. However, despite the absence of legal requirements, some UK universities currently require ethics review processes for insect research (see, e.g., Aberystwyth University, 2025). This voluntary expansion of the scope of ethics review outside of ASPA-regulated species represents an important development in animal research oversight.

This precautionary approach to ethics review for insects in research has scientific and ethical justification. Recent neurobiological and behavioural evidence suggests it is plausible that at least some insects may be sentient, understood as the capacity to experience affective states (such as hunger or pain; Gibbons et al., 2022; Crump et al., 2023; Barrett and Fischer, 2025), a view shared by the majority of surveyed animal behavior scientists (Zipple et al. 2024). Indeed, as assessed by the framework that led to the UK’s Animal Welfare (Sentience) Act 2022 recognising cephalopod molluscs and decapod crustaceans as sentient, empirical support for adult insect sentience is stronger than the support found for adult decapod crustaceans (see Birch et al., 2021 v. Gibbons et al., 2022), particularly in the Diptera and Blattodea.

As a result of the accumulating evidence, many scientific societies (e.g., Animal Behavior Society, 2024; Insect Welfare Research Society, 2023; Royal Entomological Society, 2024) and consensus documents (Low, 2012; Andrews et al. 2024) now support the plausibility of sentience in insects while recognizing the need for further research. Many of these same groups thus recommend a precautionary approach, where some steps are taken to reduce the risk of causing unnecessary harm where insects are used or managed (Birch 2017) even while acknowledging the significant uncertainty surrounding insect sentience. The precautionary approach gains additional support from UK public opinion polls that demonstrate high levels of support (91.06%) for protecting animals when scientific experts suggest potential sentience, even in the absence of definitive proof (Rethink Priorities, 2021; and see: Drinkwater et al. 2019). Proportionate measures to improve the ethical treatment of insects in research can thus protect human health and wellbeing while managing public opinions and promoting insect welfare (Freelance, 2018, Drinkwater et al. 2019).

However the expansion of ethics review to include insects at the institutional level may introduce unique challenges, both for researchers and reviewers. On the researcher side, a lack of experience can hamper successful communication with the ethics committees (Harvey-Clark, 2011), especially as surveys of entomologists reveal limited familiarity with animal welfare concepts or training on insect welfare (despite strong support for both the training and resources to improve practice; Barrett et al., 2025). More broadly, biologists receive minimal ethics training during their education, with graduate students often not being exposed to ethical frameworks applicable to their research (Trout et al. 2010; Crozier & Schulte-Hostedde, 2015). This training gap becomes particularly problematic given the distinctive challenges posed by insect research ethics. Unlike research with traditional laboratory animals, insect research spans tremendous physiological and behavioural diversity across taxa, and occurs in diverse contexts; however, there remain very few established welfare assessment tools adapted to invertebrates’ unique biology and their contexts of use (Barrett & Fischer, 2023).

On the reviewer side, there can be limitations in expertise for specialised research areas, as well as general insect biology, that make assessment difficult. While animal research ethics has long been guided by the principles of reduction, refinement, and replacement (Russell & Burch, 1959), the application of those principles looks quite different in different contexts. Animal research ethics was once focused on the use of a few species in laboratories; now, it has expanded to an enormous array of species and research contexts. As a result, scholars in some fields, such as ecological and wildlife research, have identified significant gaps in ethical guidance (even when working with vertebrates), and have called for more systematic approaches to ethical research design, particularly for non-model species and field research contexts (Crozier & Schulte-Hostedde, 2015; Field et al., 2019). Ultimately, the absence of guidance in non-standard contexts or on non-standard species makes oversight more complicated for those offering ethical review and for researchers. Such issues can be compounded by the growing workload and responsibilities of ethics review committees, which can slow down the review process and lead to dissatisfaction with review processes within research communities (Page & Nyeboer, 2017).

Despite the adoption of ethics review for insect research at some UK universities, no research has examined how wide-spread this expansion may be, how it is implemented by institutions, or how experienced by researchers. This study addresses these knowledge gaps by providing the first systematic examination of the number of UK institutions that have formal ethics review processes for insects and UK entomologists’ experiences with and attitudes toward ethics review for insect research. Through a comprehensive survey of Royal Entomological Society members and direct contact with university ethics boards, we examine researchers’ experiences with ethics review (voluntary and mandatory), their attitudes towards ethics application, and what resources and support they use to facilitate ethics applications Separately, we contacted universities and institutions to determine current ethics requirements for those working with insects.

## Materials and Methods

### Confirmation of university requirements

142 universities and institutions, identified through Royal Entomological Society membership lists (Supplemental File 1), were directly contacted through their ethics board to ascertain current ethics requirements for those working with insects. Emails were sent in October 2024 with follow up correspondence from those that did not respond in December 2024, March 2025, May 2025, June 2025 and August 2025.

### Ethics Approval, Study Design, and Participant Criteria

This survey portion of this study was reviewed and approved by the Human Behaviour Change for Life Ethics Committee (HBCL006Ins).

An online survey was designed in SurveyMonkey (California, USA) to explore the views of researchers working with insects in regard to ethics applications. The survey consisted of 22 questions (Supplemental File 2) which branched into three streams depending on if the researcher was subject to mandatory, voluntary, or no, ethics application processes for their work. The survey was piloted by 5 research scientists that work with insects across disciplines and institutions, all active members of RES. Participation was voluntary, with researchers recruited to the study through the RES newsletter, social media, and personal correspondence by RES staff only.

The survey was open to those over the age of 18, based in the UK, that self-selected as working with insects in research. Questions explored the researchers’ background, current ethics application status for insect research, experiences with ethics review for insect research (voluntary and mandatory), any resources and support used to facilitate ethics applications when using insects (voluntary and mandatory), and attitudes towards ethics review of insect research.

### Qualitative Data Coding

Qualitative data on the impact of ethics review on the quality of research and any additional comments were coded by a member of the research team with both entomological knowledge and qualitative data analysis experience using methods described in Creswell and Poth (2018). First, an exploratory analysis was conducted: responses were read several times to obtain a general sense of the dataset. Then, data were manually coded for each qualitative response. For each response, text segments were identified and linked to a single code with a description. After all coding, the set of codes was analysed for overlap and redundancy and descriptions were refined. Themes were identified by aggregating together similar codes that were about a similar idea. Finally, quotes from participants (anonymised by providing each participant with a randomly generated 3-number ID) were obtained from the dataset as representative examples to provide in the codebook (Supplemental File 3).

### Data Cleaning and Statistical Analyses

The survey for researchers was open from the 20^th^ January 2025 to the 30^th^ April 2025. All online responses were downloaded from SurveyMonkey on the 1^st^ May 2025. Responses were checked for consistency and screened for multiple responses. Any insufficiently completed responses, duplicate responses, or responses from those based outside of the UK, were removed from the master copy. Once cleaned all names were removed from the master sheet to ensure anonymity. Data were analysed in R (v4.4.2; R Core Team 2025). Due to the small sample size of voluntarily-submitted applications (n = 5), voluntary and mandatory ethics applications have been grouped for all further analyses.

## Results

### University and Institution Ethics Processes

In total of 142 UK universities and institutions were contacted, with 109 providing responses (77% response rate). Among these, 14.7% (n = 16) stated no insect research is conducted at the university and were removed from analysis. Of the remaining 93 institutions, just over half (51.6%, n = 48) reported that ethics applications were required for work with insects, while 48.4% (n=45) reported that they do not currently require ethics applications for working with insects. Notably, over a third (35.6%) of the universities and institutions that do not currently require ethics applications expressed interest in being informed of resources produced on the ethical treatment of insects in research.

The Royal Entomological Society maintains a list of universities considered to have an entomological course. This list is maintained internally through searches and correspondence with course leaders. In 2025, there were 46 universities on this list. Of these universities, 37 responded (80%). 56.8% (n = 21) do not require ethics applications for working with insects, while 43.2% (n = 16) do require ethics applications.

### Survey Response and Demographics

In total 145 respondents participated in the survey. Following data cleaning, 129 responses remained for analysis (Figure 1).

**Figure 1:**
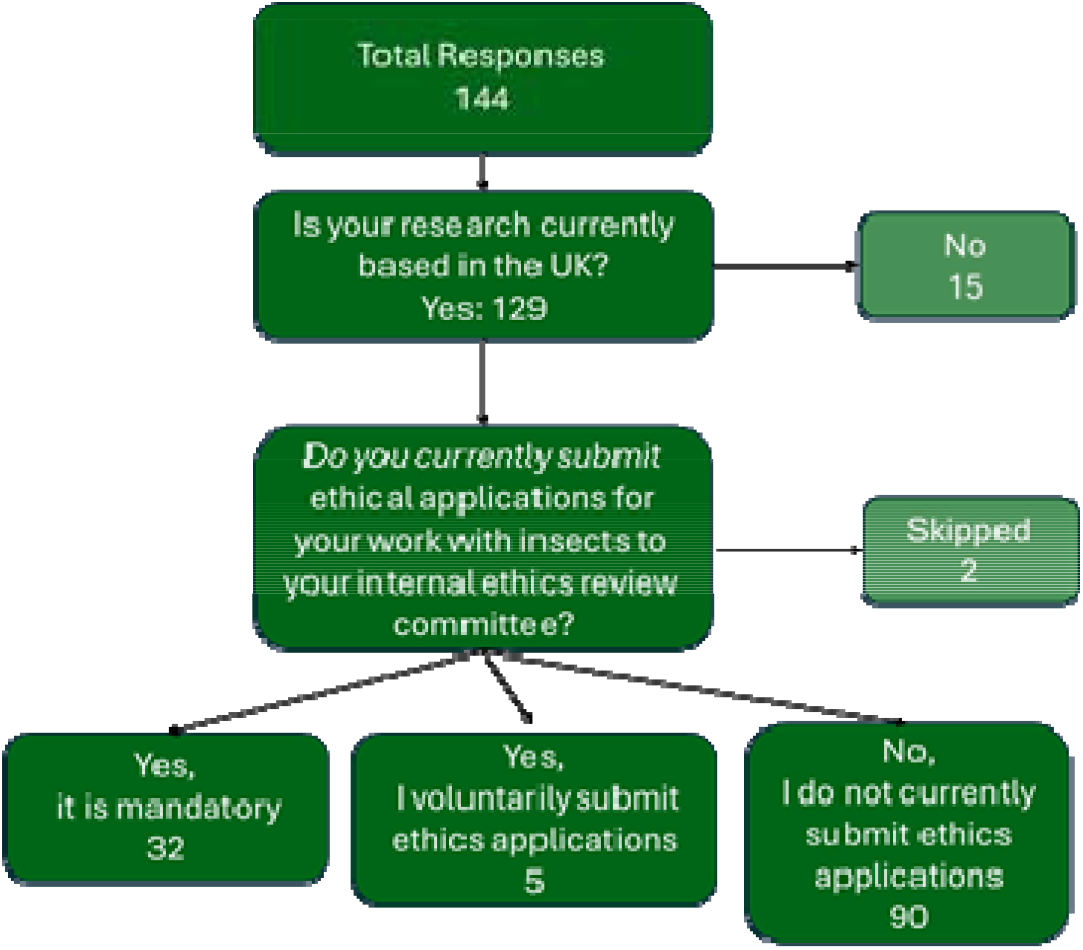
Survey response rate.

Respondents spanned career stages and primary areas of research (Table 1). A total of 53 named institutions were represented, of which only 14 universities/institutes represented ≥ 3 responses.

**Table 1.**
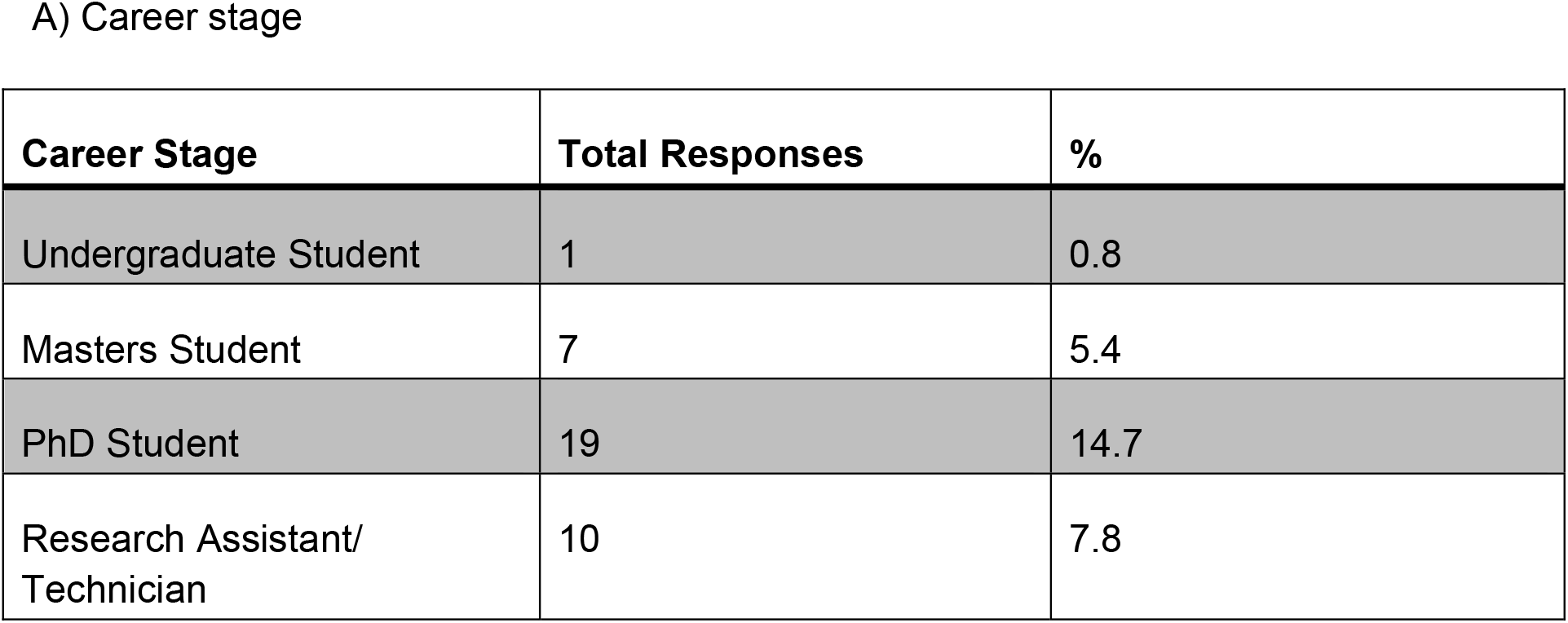
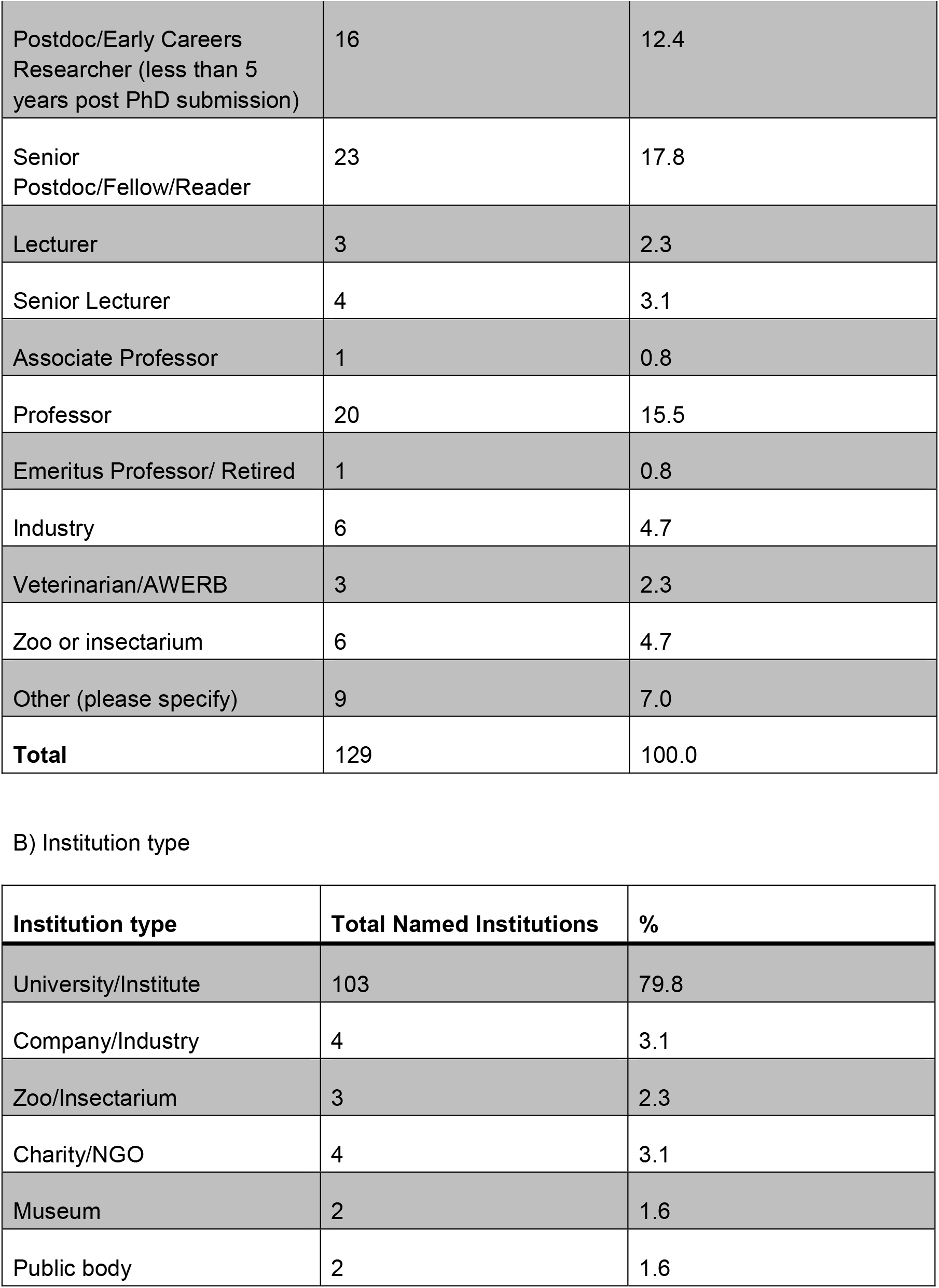
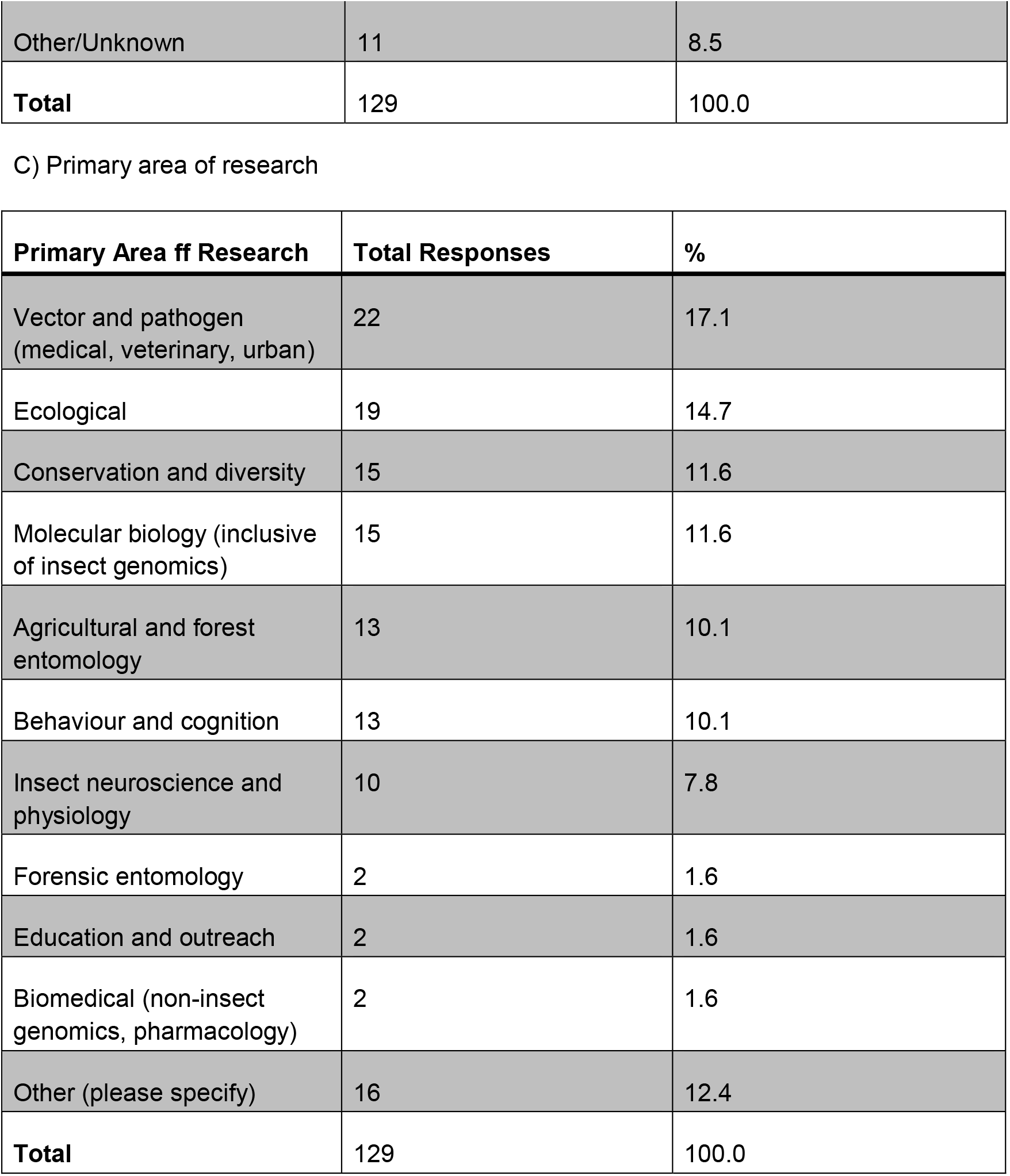
Respondent demographics.

A) Career stage
| Career Stage | Total Responses | % |
| --- | --- | --- |
| Undergraduate Student | 1 | 0.8 |
| Masters Student | 7 | 5.4 |
| PhD Student | 19 | 14.7 |
| Research Assistant/<br>Technician | 10 | 7.8 |
| Postdoc/Early Careers Researcher (less than 5 years post PhD submission) | 16 | 12.4 |
| Senior Postdoc/Fellow/Reader | 23 | 17.8 |
| Lecturer | 3 | 2.3 |
| Senior Lecturer | 4 | 3.1 |
| Associate Professor | 1 | 0.8 |
| Professor | 20 | 15.5 |
| Emeritus Professor/ Retired | 1 | 0.8 |
| Industry | 6 | 4.7 |
| Veterinarian/AWERB | 3 | 2.3 |
| Zoo or insectarium | 6 | 4.7 |
| Other (please specify) | 9 | 7.0 |
| **Total** | **129** | **100.0** |

B) Institution type
| **Institution type** | **Total Named Institutions** | **%** |
| --- | --- | --- |
| University/Institute | 103 | 79.8 |
| Company/Industry | 4 | 3.1 |
| Zoo/Insectarium | 3 | 2.3 |
| Charity/NGO | 4 | 3.1 |
| Museum | 2 | 1.6 |
| Public body | 2 | 1.6 |
| Other/Unknown | 11 | 8.5 |
| **Total** | 129 | 100.0 |

C) Primary area of research
| Primary Area of Research | Total Responses | % |
| --- | --- | --- |
| Vector and pathogen<br>(medical, veterinary, urban) | 22 | 17.1 |
| Ecological | 19 | 14.7 |
| Conservation and diversity | 15 | 11.6 |
| Molecular biology (inclusive<br>of insect genomics) | 15 | 11.6 |
| Agricultural and forest<br>entomology | 13 | 10.1 |
| Behaviour and cognition | 13 | 10.1 |
| Insect neuroscience and<br>physiology | 10 | 7.8 |
| Forensic entomology | 2 | 1.6 |
| Education and outreach | 2 | 1.6 |
| Biomedical (non-insect<br>genomics, pharmacology) | 2 | 1.6 |
| Other (please specify) | 16 | 12.4 |
| **Total** | 129 | 100.0 |

Entries are n (%). UK respondents defined as ‘Yes’ to “Is your research currently based in the UK?”. A comparison of RES membership by career stage (student, member, retired member) can be found in Supplemental Table 4.

70.9% of respondents (n = 90) reported that they did not currently submit ethics applications for their work with insects. 25.2% (n = 32) reported mandatory ethics applications for their work, whilst 3.9% (n = 5) choose to voluntarily submit ethics applications for their work. Manual comparisons between the university-reported data on ethics processes and researcher, self-reported ethics applications highlighted 7 researchers, from 5 different institutions, that believed they did not need to submit ethics applications for their work with insects despite their institutions stating that work with insects would fall under their ethics review process.

The earliest researcher report of mandatory ethics application for working with insects was 2008 (Figure 2). Among researchers that submitted ethics applications in the last year, the majority (51.4%) submitted just 1-2 applications, with 7/35 (20.0%) submitting none; 2/35 (5.7%) reported 6–10 applications and 2/35 (5.7%) reported 11–20 applications.

**Figure 2:**
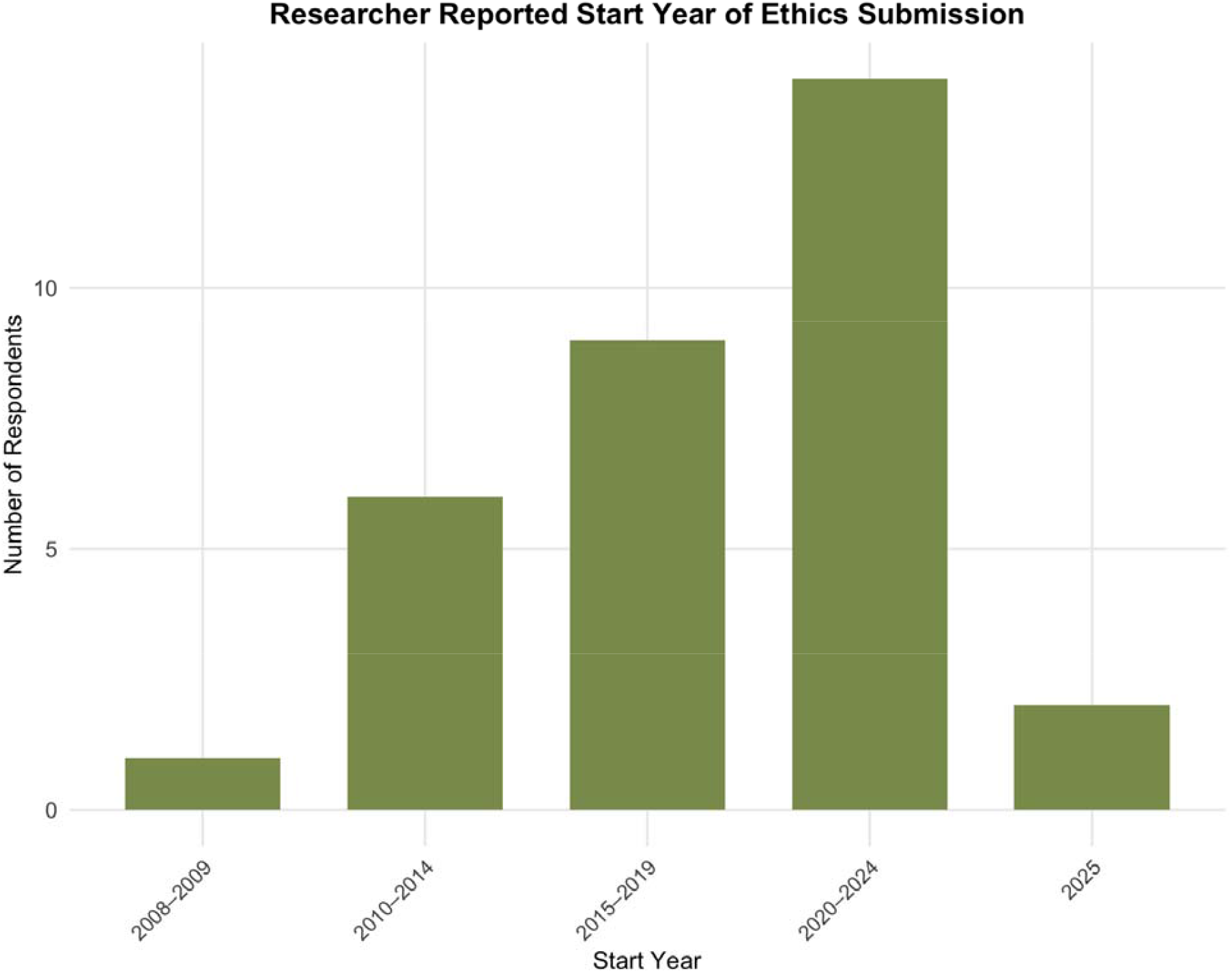
The earliest report by researchers of mandatory ethics review applications for working with insects.

### Reasons for submission and feedback received

In a ‘select all that apply’ question, 73% of researchers that submitted ethics applications reported institutional requirement as the reason for submitting (n = 27/37). Other commonly reported reasons included personal ethical beliefs (32%, n = 12), ensuring subject welfare (32%, n = 12), and scientific best practice/standardisation (32%, n = 12). Less frequently reported reasons were grant requirements (13.5%, n = 5), requirement to publish (10.8%, n = 4), to receive panel feedback (5.4%, n = 2) and public perception (2.7%, n = 1).

In a ‘select all that apply’ question, 43% indicated they received no feedback from the ethics review board on their application (n = 15, plus one researcher that answered ‘other’ but then indicated acceptance with no feedback in the comment box). 24% (n = 9) of researchers selected ‘positive comments’, 19% (n = 7) of researchers selected ‘rejections with reasons provided’ (with one researcher commenting that resubmission was possible to address the reasons provided in the comment box), and 14% (n = 5) of researchers selecting ‘suggestions for improvement of study methodology’. In the comment box, examples of feedback received included: requests for more information on the methodology, better justifications of sample size, and better justification or consideration of refinements including considerations for anesthesia, disease management, enrichment, and/or euthanasia.

### Confidence in ethics applications

Self-reported confidence amongst those that submitted ethics was moderate to high in both the ethics application process and ability to deliver what is expected when submitting an ethics application (Table 2). There was no discernible difference in confidence in the ethics application process across years of experience (n=32, Kruskal-Wallis χ ^2^(5)= 3.45, p=0.63).

**Table 2.** Confidence in the ethics application process (1) and ability to deliver what is expected (2) when submitting an ethics application amongst those reporting ethics submissions for their work with insects.

| Self reported confidence | (1)"How confident do you feel about the ethics application process?"<br>(n=32)<br>(%) | (2)"How confident are you that you can deliver what is expected of you when completing an ethics application?" (n= 32)<br>(%) |
| --- | --- | --- |
| Not confident at all | 9.4 | 3.1 |
| Somewhat not confident | 0.0 | 0 |
| Neutral | 21.9 | 15.6 |
| Somewhat confident | 46.9 | 34.4 |
| Very Confident | 21.9 | 46.9 |

**Table 2.**
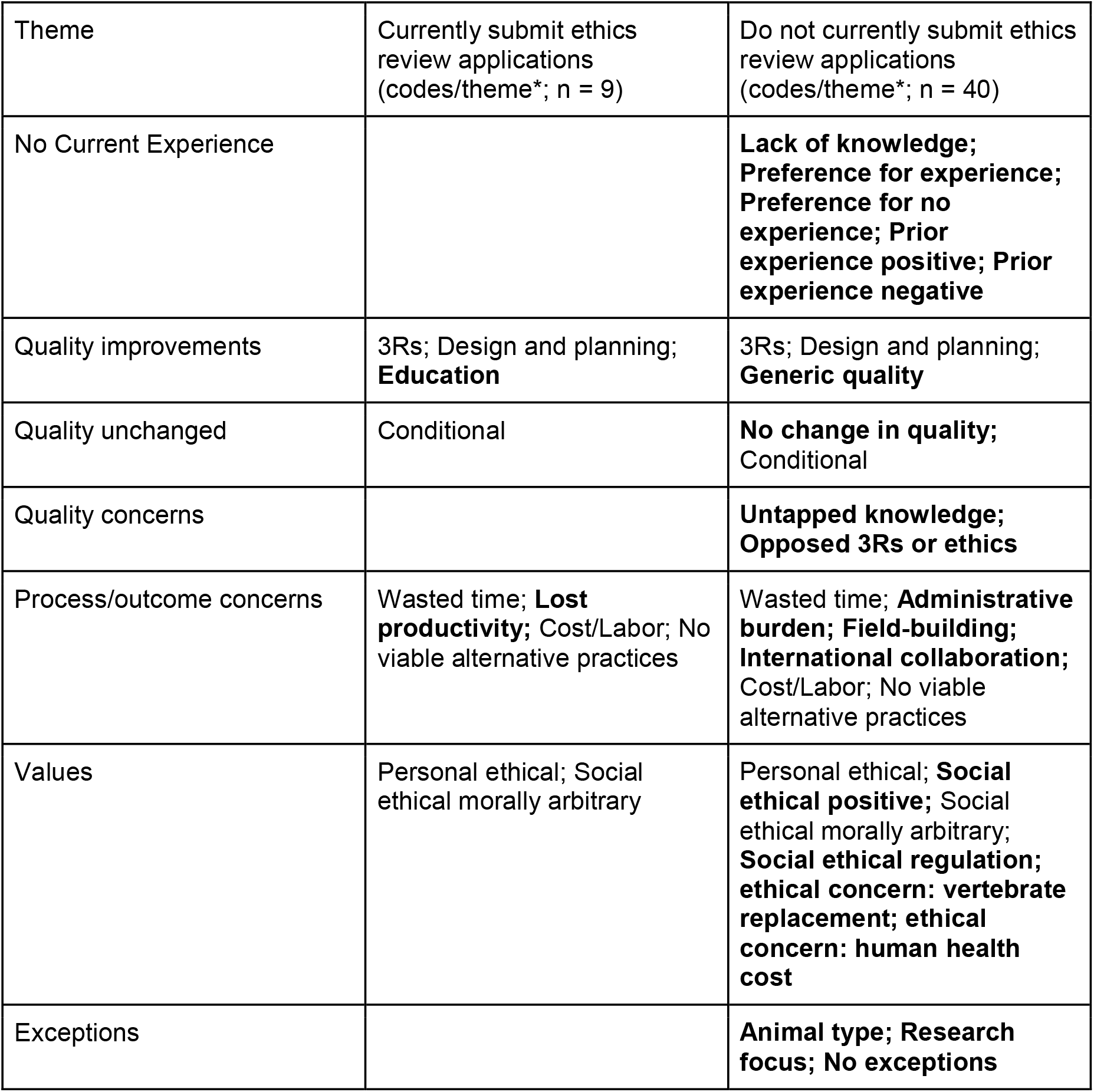
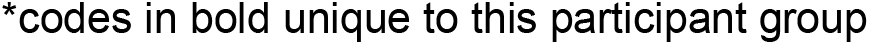
Themes and codes that emerged from qualitative analysis of participant responses to the question “In your opinion, how does the ethics application process influence the quality of research?” (Q1)

| Theme | Currently submit ethics review applications<br>(codes/theme*; n = 9) | Do not currently submit ethics review applications<br>(codes/theme*; n = 40) |
| --- | --- | --- |
| No Current Experience |  | **Lack of knowledge; Preference for experience; Preference for no experience; Prior experience positive; Prior experience negative** |
| Quality improvements | 3Rs; Design and planning; **Education** | 3Rs; Design and planning; **Generic quality** |
| Quality unchanged | Conditional | **No change in quality;**<br>Conditional |
| Quality concerns |  | **Untapped knowledge; Opposed 3Rs or ethics** |
| Process/outcome concerns | Wasted time; **Lost productivity;** Cost/Labor; No viable alternative practices | Wasted time; **Administrative burden; Field-building; International collaboration;** Cost/Labor; No viable alternative practices |
| Values | Personal ethical; Social ethical morally arbitrary | Personal ethical; **Social ethical positive;** Social ethical morally arbitrary; **Social ethical regulation; ethical concern: vertebrate replacement; ethical concern: human health cost** |
| Exceptions |  | **Animal type; Research focus; No exceptions** |

\*codes in bold unique to this participant group

Questions were optional for those that reported mandatory and voluntary ethics applications for their work with insects (n= 32 for both questions).

### Resources to support ethics applications

In a select all that apply question for those currently submitting ethics applications (“What resources or support do you think would help you (and/or your students) with ethics applications submissions”), 60% (n = 18/30) entomologists selected ‘Greater research on insect welfare’, 57% (n = 17) selected ‘training on the ethical use of animals in research with examples of how to apply this to insects’, 43% (n = 13) selected both ‘guidance on how to develop an ethical application for submission to ethics panels (AWERBs)’ and ‘information on insect biology for those on the ethics panels (AWERB members), and 40% (n = 12) selected ‘undergraduate curricular materials on insect welfare’. ‘Other’ was selected by 20% (n = 6); however, 4 responses did not answer the prompt by offering any information about resources or support for ethics applications. Of the other responses, one echoed the need for guidance on how to submit ethics applications on insects specifically and the other asked for training specific to sample sizes and experimental design.

When asked what resources were currently used, only four responses (n = 13%) were received, suggesting the majority of respondents do not use any resources when crafting ethics applications. One entomologist mentioned the ASAB ethics guidance for nonhuman animals, the NC3Rs website and Home Office legislation, while another mentioned the Veterinary Invertebrate Society, a third mentioned ‘papers’, and the fourth mentioned their undergraduate course.

### Perception of ethics on research quality

Perceptions of the influence of ethics on research quality did not differ by submission status (χ^2^(2) = 0.31, p = 0.86; Cramer’s V = 0.06); therefore, all respondents (n = 101) were analysed together.

Perceptions of the influence of ethics applications on research quality were mostly positive. Across all respondents, 44.6% (n = 45/101) reported improvements in research quality as a result of ethics applications, 39.6% reported no effect, and 15.8% reported that ethics applications detracted from research quality. When restricting analysis to evaluative responses (e.g., excluding ‘no effect’), 73.8% judged that ethics review improves quality (95% CI 60.9– 84.2; exact binomial p<0.001; Figure 3). As there was no difference in perception between those currently submitting and not currently submitting, this suggests that sentiment is broadly favourable across respondents, rather than just confined to those currently submitting ethics for their work with insects.

**Figure 3.**
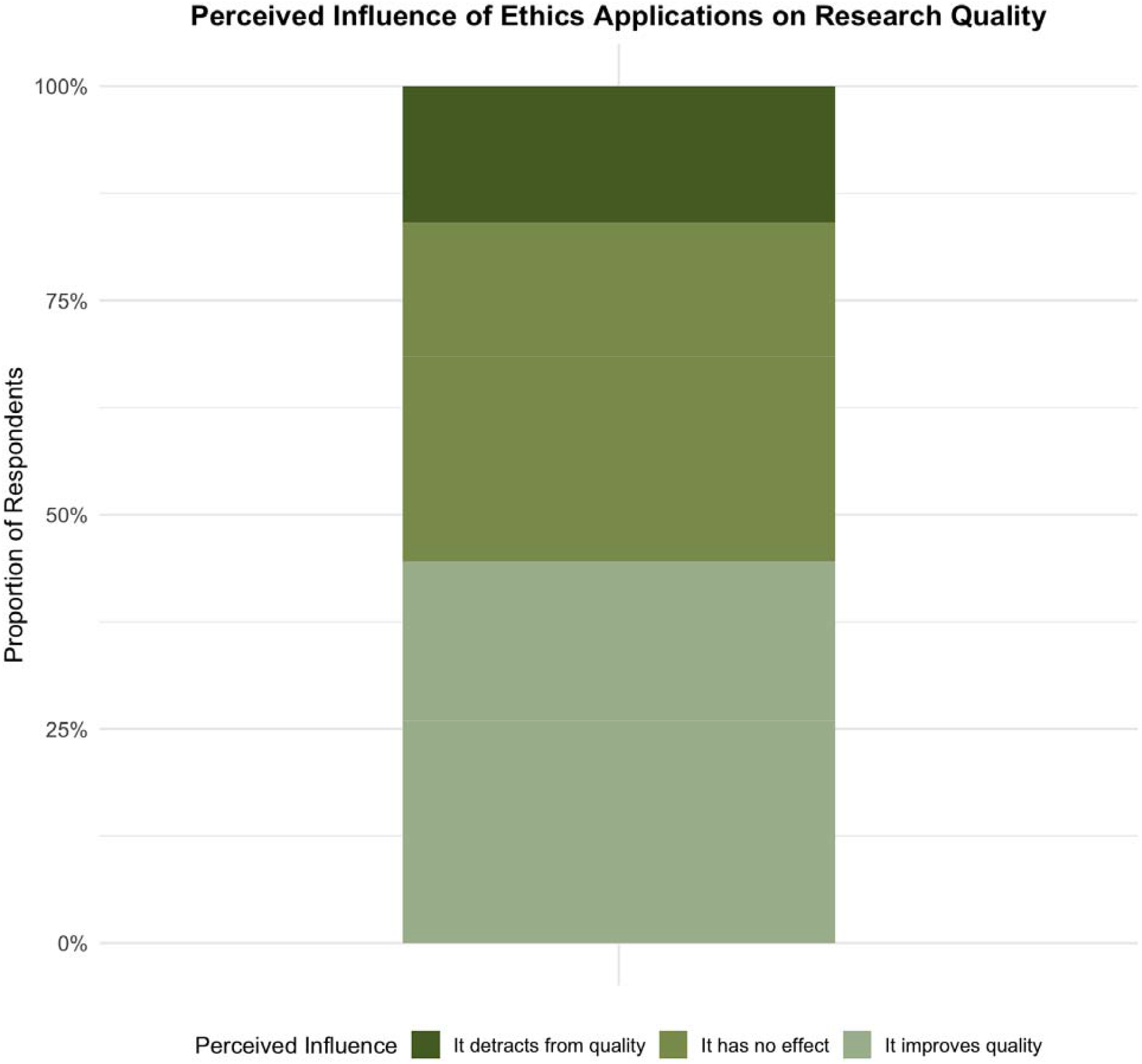
Opinion on the influence of ethics on quality (voluntarily submit ethics applications (n= 3), mandatorily submit ethics applications (n=28) and those that do not submit ethics applications (n=70)).

Due to small group size, those reporting <1 year experience working with insects (n = 3) were grouped with 1-2 years experience (n = 14, combined ≤2 year group n = 17) for analysis. The perceived influence of ethics on quality was inversely correlated to years of experience, with those newer to research stating ethics applications improved quality (Figure 4) (70.6% ≤2 years, 52.9% 3-5 years, 47.4% 6-10 years, 47.4% 11-20 years, 20.7% ≥21 years). A linear trend test was significant (χ^2^(1)= 10.56, p=<0.001) and in a binomial model each higher band (e.g., increased years of experience) was associated with lower odds of stating ‘improves’ (OR per band 0.62, 95% CI 0.46-0.83).

**Figure 4.**
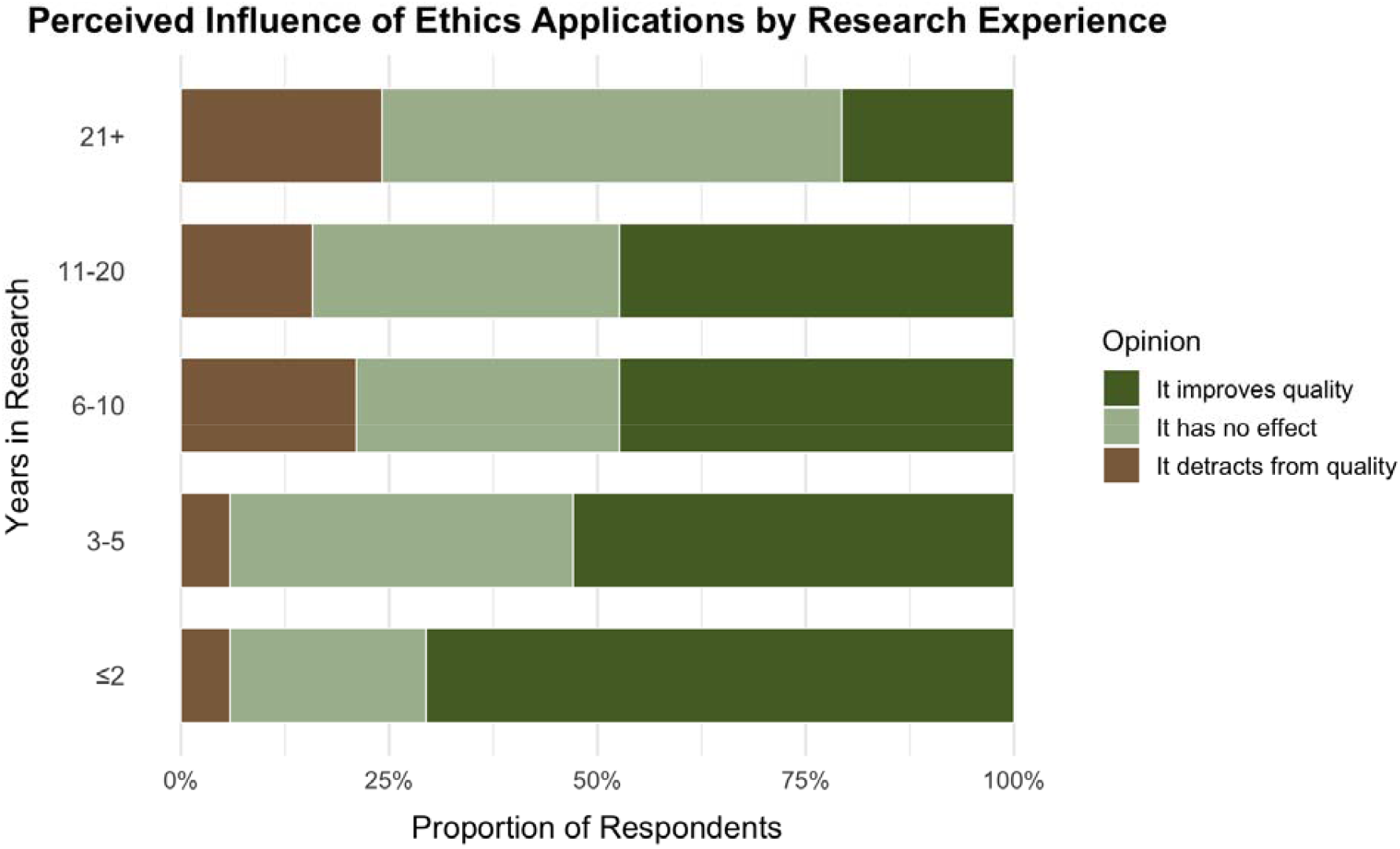
Opinion on the influence of ethics on quality by years of experience: 21+years (n=29), 11-20 years (n=19), 6-10 years (n= 19), 3-5 years (n=17), grouped <2 years (1-2 years (n=14) and <1year (n=3)).

Of those that submit ethics applications (n = 31), only 16.1% reported changes to their practices attributed to the ethics process, with 74.2% reporting no change (9.7%, unsure). In the qualitative responses asking for participants to explain any changes in practice, those that reported improved research quality from ethics applications noted that they made changes to species collection protocols, generically improved experimental design, or reduced their lethal trapping (n = 3 qualitative responses). Those that reported their practices had changed as a result of ethics applications in a way that detracted from research quality noted that they now needed to complete additional paperwork and that, although they had not yet changed this practice, they were considering switching to vertebrates because of ethics review for insects (n = 2 qualitative responses).

All participants (irrespective of submission status) were then asked the free-response question “In your opinion, how does the ethics application process influence the quality of research?”. Some themes and codes that emerged were shared irrespective of whether participants currently submitted ethics review or did not currently submit ethics review (Table 2; Supplemental File 3), but some were unique to each group (though sample sizes for qualitative responses were limited).

Those that did not currently submit ethics applications often noted their lack of current experience with ethics review, though some described prior positive or negative experiences with ethics review for insect research. Additionally, some described a preference to submit ethics applications (“I have no experience in submitting an ethics application. In retrospect I wish I had the opportunity” [P811]) or a preference not to submit ethics applications (“I… would prefer not to see this instituted for the kind of work I do” [P744]).

When it came to the substantive question of research quality impacts, some participants from both groups indicated that research did (currently submitting) or was predicted to (not currently submitting) improve in quality as a result of ethics application, as a specific result of the 3Rs or increased time spent on study planning: “Particularly regarding ‘reduction’, there is more thought put into making the most out of data and sampling efficiently in ecological studies” [currently submits; P391] or “I can see how the ethics application process would make researchers more mindful of experimental design etc by having to consider correct use[s]age of animals and the correct controls” [does not currently submit; P384]. One participant that currently submits ethics applications described ethics review as an opportunity to educate researchers on quality-enhancing components of experimental design (“ethics application[s]… educate researchers on power calculations and emphasize the key objectives of research” [P175]). Only members of the group that did not currently submit described generic endorsements of research quality improvements from ethics review, compared to identifying specific aspects of ethics review that caused improvements in research quality.

Concerns about direct, negative impacts on research quality were also only raised in the qualitative responses by participants not currently submitting ethics review. One participant described their belief that the 3Rs were antithetical to research quality: “The ethics application process detracts from the quality of research…There are many areas of insect research in which Replacement, Reduction, and Refinement are antithetical to the research itself” [P204] before providing some examples where there were no alternative practices (another code) available in their discipline. More commonly, researchers not currently submitting ethics review expressed the concern that some research areas may become off-limits, thereby reducing scientific understanding or applied knowledge: “It has an effect on the things people can research, putting a limit to certain branches of experimentation” [P682]. Others not currently submitting predicted that ethics review was not expected to change study quality: “their quality of work will stay the same” [not currently submitting; P682].

Finally, some participants from each group indicated that quality changes (positive or negative) could be conditional: “If done well, it does not have to detract from research quality” [currently submitting; P130] or “In my opinion, however, whether going through ethical approval could have a positive or negative effect on the research may depend on purpose and target group.” [not currently submitting; P629]. These conditional responses to ethics review of insect research were also exhibited in response to the final question on the survey.

Beyond research quality concerns, participants also expressed concerns with the way ethics review could impact scientific processes or outcomes (some of which could indirectly impact quality), a variety of ethical values or concerns, and possible exceptions (or lack of exceptions) to ethics review. As these do not relate to research quality *per se*, and were also exhibited in responses to the final general comment question covered next, we will cover these kinds of responses to both this question and the following in the subsequent section.

### General qualitative insights on the ethical treatment of insects in research

The last question of the survey asked all participants to write a response to: “Please share any additional comments or insights regarding your experience with, or attitudes about, the ethics application process in entomological research”. Many participants seemed to strongly link their responses to the prior question on research quality to this general response question (‘linked responses’, “see previous comment” [P685]), resulting in many similar codes and themes emerging in the qualitative analysis of each question (Table 3; Supplemental File 3).

**Table 3.**
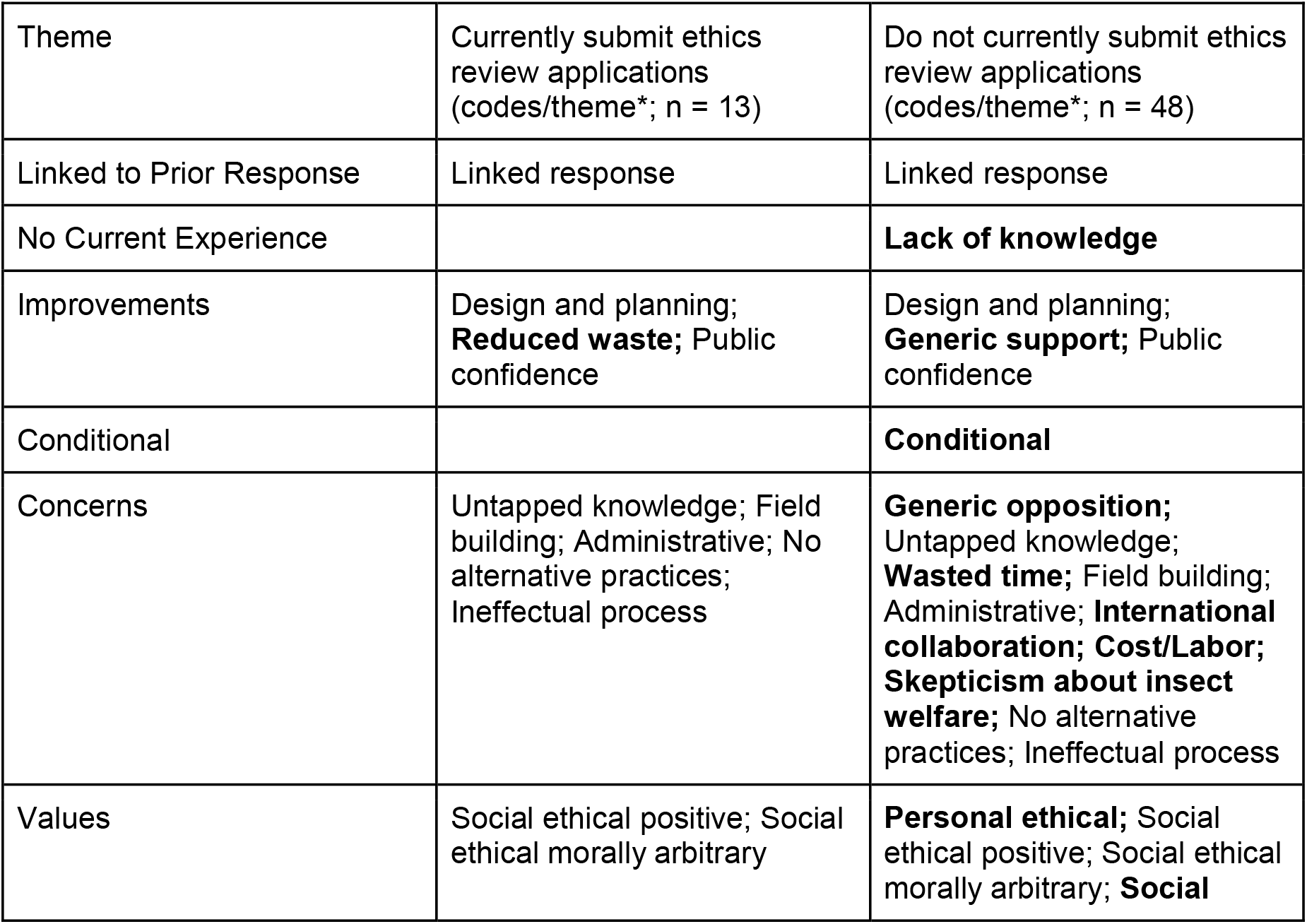
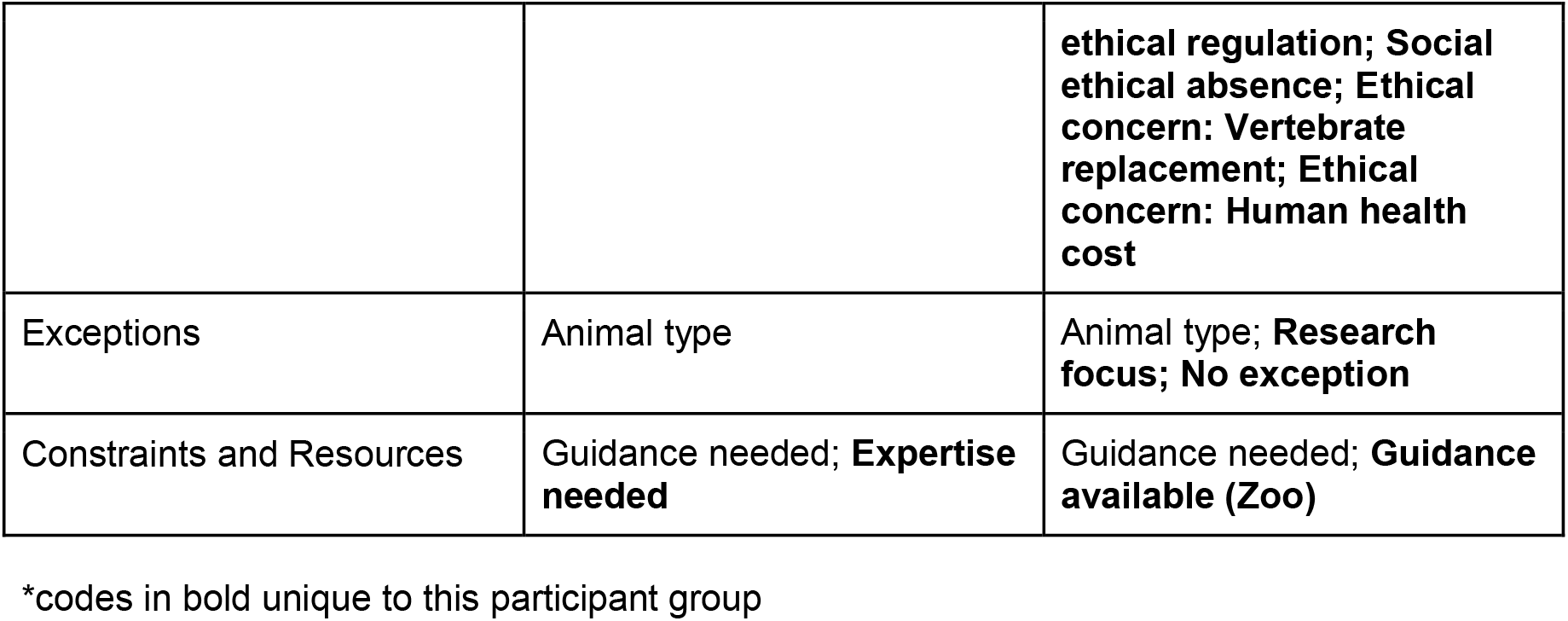
Themes and codes that emerged from qualitative analysis of participant responses to the question “Please share any additional comments or insights regarding your experience with, or attitudes about, the ethics application process in entomological research.” (Q2)

Participants of both groups described additional reasons for supporting ethical review, reiterating design and planning improvements as a result of ethical consideration, describing reduced waste that had occurred as a result of ethics review (“In principle this has forced some research to change how they approach research, where they used to have huge animal (insect wastage.” [currently submits; P389]), discussing positive impacts on public confidence in science (“It is/should be an integral component of the research process and can improve the integrity, transparency, public confidence and the design and quality of research” [currently submits; P605]), or offering generic supportive comments in the case of those not currently submitting (“I think this is a great topic to raise and discuss” [not currently submitting; P502]).

Alongside generic support, there was also generic opposition to ethics review by those not currently submitting: “I do not feel that it is valuable to apply ethics to insect experimentation” [P238] or, more strongly, “Applying ethics would significantly damage research” [P498]. As in the prior question, some respondents reiterated that improvements to research as a result of ethics review were not guaranteed, but rather conditional on a variety of ways in which the process could be or was applied.

However, many participants from both groups described specific concerns with ethics applications in either this question [Q2] or the prior question [Q1], including a variety of research quality and process/outcome concerns:

- Untapped knowledge: “such a stance could inadvertently hinder broader entomological research. While working with insects for human health and agricultural purposes will likely remain widely accepted, introducing ethical ambiguity could impose unnecessary restrictions on other areas of study” [not currently submitting, Q2; P798];
- Wasted time: “A huge amount of time is being wasted on ethics procedures” [currently submitting, Q1; P728]
- Increased cost/labor: “having a committee solely focused on this would no doubt improve quality, but at financial cost” [not currently submitting, Q1; P712]
- No viable alternative practices: “I work with Drosophila melanogaster…so I have hundreds of strains that I can only maintain by live propagation (no freezing), so I have no other solution that raising and killing millions of flies every year” [currently submitting, Q1; P189]
- Increased administrative burden: “it would add unnecessary bureaucracy to an already very bureaucratic existence” [not currently submitting; P744]
- Negative impacts on field building: “I fear this would make it more difficult to carry out research, particularly in a research field where we need to be encouraging more people to join” [not currently submitting, Q2; P293]
- Ineffectual process: “I feel like it is usually always approved as it is more a box ticking exercise” [currently submitting, Q2; P562]; “often the suggestions are not very helpful” [currently submitting, Q2; P736]; “They’re usually incredibly lack-lustre and do little to prevent neglect of a species” [currently submitting, Q2; P407]

One scientist currently submitting ethics applications described a realised concern with lost productivity: “blanket imposition of burdensome regulations has already impacted research productivity” [Q1; P130]. Only among those not currently submitting, researchers expressed concerns that international collaboration could be made more difficult (“[ethics applications would] make it more difficult to collaborate with countries that do not impose such restrictions” [not currently submitting, Q2; P593] or “Adding these ethics regulations will directly harm efforts to build equality in global health” [not currently submitting, Q1; P204]).

Skepticism that insects either had sentience/welfare at all, or that their welfare could be robustly described using empirical tools, were not found in the subset of responses from individuals currently submitting ethics applications, but were provided by several respondents in the group of scientists not currently submitting, for instance:

- “The attitude of researchers and ethical guidelines to our practices should reflect the knowledge that insects feel pain, and that we are uncertain of whether or not they experience suffering” [not currently submitting, Q2; P932];
- “insects live a very different existence. There is no evidence that they experience pain as we do” [P498]
- “Currently we have a debate in the field as to whether insects are sentient or not. In our lab, we take the stance that insects may be sentient, but there is no scientific evidence to prove this, and therefore are not subject to consideration for welfare and ethics from this perspective… [a shift to discussing research outcomes instead of insect welfare] will prevent the discussion being led by subjective and ill-defined concepts of what insects may feel or may not feel” [not currently submitting, Q2; P210]

Those that were not currently submitting ethics review expressed concern in response to these questions that ethics review for insects could also lead to ethical problems, including reduced replacement of vertebrates in research (“It would also increase use of vertebrates due to the ‘insect advantage’ of no regulatory ethics being removed” [not currently submitting, P498]) or human health costs (“If the RES were to misinterpret or overstate the ethical considerations surrounding insect sentience, it could have significant consequences for research critical to human health, particularly in vector biology and the use of Drosophila as a model organism” [not currently submitting, Q2; P798]; or “If you [those conducting research in this area] continue, you will kill human beings” [not currently submitting, Q2; P204]).

Some participants noted that ethics review would require better practical guidance on insect welfare; those from the zoological community noted that they already had guidance from their professional organization (BIAZA) on terrestrial invertebrates. Additionally, one researcher currently submitting applications noted that subject area/insect expertise was missing on their ethics review panel and that “Making sure that the correct people are on the panels, especially when this is expanding, we need to make sure we have capacity and representation” [Q2, P736].

Some participants from both groups described ways they believed exceptions for specific animals or specific types of research were necessary or could mitigate perceived concerns with ethics application. Generally, this was their own study organism or area of research:

- “I would very much object if someone came to my lab to tell me what to do with them [millions of Drosophila]…A different matter would be if I worked with stag beetles; then, I would really like to have guidelines about what others find objectionable, besides my own considerations. But luckily I don’t and never will. I would find it reasonable to question whether any planned research that harms stag beetles needs to be carried out at all, and whether the same knowledge cannot be obtained from another species or in another way” [currently submitting, Q1; P189]
- “For example, if research is done on an endangered insect for a conservation process, there could be benefits from considering ethics of procedures. In my case, where research focussing on understanding ecology of pest/vector species that proliferate and can cause huge impacts on human and animal health, having to go through an extensive ethical approval process… would be a detraction” [not currently submitting, Q1; P629]

Many participants emphasised that insect research should be treated differently from vertebrate/mammal research when it came to ethics review: “Separation of vertebrates & invertebrates made clear for ethics approval requirements” [not currently submitting, Q2; P436]. However, there were also researchers that opposed exceptions for specific animals, including participants who believed the same standard of ethics review should be used for vertebrates and insects (“I believe that there should be thorough ethical assessment for invertebrate work, and that it should be similar to ethics applications for working with vertebrates” [not currently submitting, Q2; P141]). Some researchers also believed the same standard of ethics review (or lack thereof) should be used for all areas of insect research, including from a participant opposed to ethics review entirely: “I oppose arguments that vector research is different than curiosity driven research [when it comes to ethics applications]” [not currently submitting, Q1; P204].

Finally, many described personal ethical views on the treatment of insects in research (some in favor of welfare considerations, others not in favor) or a variety of socially-situated views on the ethical treatment of insects in research, including:

- a perceived moral arbitrariness between the treatment of insects in research and in the world/wild (‘social ethical morally arbitrary’): “There are more insects killed and becoming extinct because of habitat loss and pesticides use than on research.” [not currently submitting, Q1; P685]
- regulation-based ethical views (‘social ethical regulation’): “not applicable, Drosophila is not a protected species under Animal act 1986 so does not require ethical approval…[so under ASPA] they are not considered animal” [not currently submitting, Q1; P468]
- a social environment that encouraged consideration of insect welfare in research as a shared cultural value and practice (‘social ethical positive’): “I support [instition]’s idea around sanctity of life and trying therefore to no[t] cause harm wherever possible” [currently submitting Q1, P389] or “we always submitted ethical review proposals for all invertebrate work and they were reviewed and discussed by all staff (it was a small biotech) and only went ahead with 100% agreement that the work was needed and that the benefit outweighed any harm” [currently submitting, Q1; P540]
- a social environment where reinforcing cultural or professional values about the consideration of insect welfare in research were absent (‘social ethical absence’): “I have never been asked by journals I publish in” [not currently submitting, Q2; P658]

## Discussion

This survey provides the first systematic overview of UK entomologists’ experiences with ethics review for insect research. Responses included individuals across career stages, institutions, and subfields, reflecting the diverse community of insect researchers. Nonetheless, the overall sample was skewed toward those not currently submitting applications, despite roughly half of UK institutions requiring them. This gap suggests that researchers subject to mandatory review may have been somewhat underrepresented. Students were also slightly underrepresented in our survey (20.9%) relative to RES membership rosters (24.5%). Future work should aim for higher engagement from these groups to provide a more balanced perspective.

Our data suggest that mandatory ethics processes for insect research in the UK have become more common over time, with the first reported institutions adopting this practice in 2008 and most adoptions coming in recent years. This growth aligns with broader precautionary trends in animal research oversight and reporting (Perl et al. 2025), as exemplified by the recent extension of protections to decapod crustaceans and cephalopods under the UK’s Animal Welfare (Sentience) Act 2022. Altogether, over half of institutions in our survey reported that ethics applications were required for work with insects. However, some researchers are not aware of their institution’s policies (as 7 researchers from 5 institutions indicated that submissions were not required when, in fact, their institution indicated that submissions are required). Greater institutional communication may be necessary if institutions want to ensure their policies are applied uniformly, especially as institutional requirement was the largest driver of ethics submissions among respondents (73%) followed by personal ethical beliefs, ensuring subject welfare, and scientific best practice/standardisation (32%, all; and see: Barrett et al. 2025).

Perceptions of the impact of ethics applications on research quality were broadly positive. Nearly half of responses indicated that ethics applications improve research quality and less than 15% indicated that ethics applications detract from research quality. Those that noted quality improved tended to focus on 3Rs and design and planning improvements. One respondent observed that “application of the 3Rs to invertebrate research has improved the design of my experiments” [P121] while another stated that “there is more thought put into making the most out of data and sampling efficiently” [P391], underscoring the constructive role that review can play in encouraging reflection and methodological rigour. Even when practices remained unchanged, participants often expressed appreciation for the reflective process itself, suggesting that ethics review may play a preventative rather than corrective role. It can ensure that welfare and methodological considerations are addressed early so that the resulting submission already aligns with best standards. Earlier career entomologists were most likely to report improved quality as a result of review (at nearly 71% for those with 2 or fewer years of experience, compared to 21% for those with 21+ years of experience). This could be because younger researchers are more open to integrating ethical reflection into their work (and see: Barrett et al. 2025) or because they benefit disproportionately from the structured planning imposed by applications. Ethics review and greater consideration of animal welfare has already been discussed as a way to address translatability and reproducibility crises in vertebrate research (Pritt and Hammer 2017; Loss et al. 2021). It is plausible, therefore, that it could play a similar role in invertebrate research, enhancing methodological rigor and promoting consistent reporting.

The majority of those that believed ethics review detracted from study quality tended to focus on the potential for knowledge to be untapped if research was not allowed to proceed due to ethics review. For instance, one participant said “It has an effect on the things people can research, putting a limit to certain branches of experimentation” [P682]. However, only 16% of researchers reported any practice changes as a result of ethics review and only 19% indicated any of their proposals had ever been rejected (with reasons provided, and some commenting that resubmission was possible). By contrast, 76% reported only positive comments, no feedback, or improvements to study methodology as a result of review, suggesting that the vast majority of research is encouraged or permitted. This matches what has been found from vertebrate animal experimentation: assessments of review committees have found they generally do not reliably require appropriate justification for animal use and that protocol approval rates are very high (Varga 2013; Schuppli, 2011; Budda et al. 2020; Milford et al. 2025; Jörgensen et al. 2020). (Indeed, some committee members self-report serving to ensure ethics review does not obstruct research objectives; see Silverman et al. 2012.) Altogether, the data from our survey and the literature on vertebrate ethics review do not suggest that ethics committees broadly act to obstruct research progress. However, developing resources that make feedback more meaningful may assist in facilitating study quality and ethics improvements as a result of review, while continuing to foster scientific progress.

Instead of quality concerns, participants were often focused on process or outcome concerns associated with ethics review, with a large majority of those being focused on lost productivity, wasted time, and increased administrative burden. This echoes concerns of scientists with other forms of ethics review, including ethics review for human research, where delays, administrative burden, and lost productivity are commonly cited complaints (Page and Nyeboer, 2017). As one challenge with ethics review is the volume of applications received by committees, adding additional classes of organisms (e.g., insects) to the workload of review committees could worsen the turnaround time of all applications without additional resources or support dedicated to review. Although this reason does not obviously justify the lack of ethics review for research subjects, it does justify institutional investment in adequate review capacity, as well as more general thought to how both ethics boards and entomologists could be supported to reduce lost productivity, wasted time, and administrative burden. To address such issues, professional societies such as the Royal Entomological Society are beginning to produce resources— including guidance documents, video materials, and trainings (as requested by some participants of this survey). These resources can support researchers in generating applications more quickly and help ethics panels streamline the application process and evaluate applications more consistently (especially where consistency and expertise are lacking: Jörgensen et al. 2020).

Another concern that some survey participants expressed was the potential negative effect of ethics review on field-building, where some believed that entomologists would be driven from the UK as a result of the burden of ethics review. However, younger entomologists were disproportionately likely to believe ethics review enhanced study quality in our survey and, in another survey, were more likely to believe training on insect welfare was important (Barrett et al. 2025). These data suggest that it is unlikely the younger generation of entomologists will be driven out of the UK as a result of ethics review. However, further research could confirm the extent of any negative effects of ethics review on field-building in entomology—as well as investigate if previously instituted animal ethics review resulted in reduced animal researchers, research productivity, and innovation in the UK.

Finally, some researchers were concerned that ethics review for insects could increase the use of vertebrate animal models in research, contradicting 3Rs efforts insofar as the 3Rs excludes invertebrates. There is no evidence that this has occurred to date as no surveyed researchers indicated they had engaged in this switch as a result of undergoing ethics review, nor does it seem particularly likely to occur frequently, given widely-accepted assumptions about which species are suitable replacements for others. (Ethical review for mouse research does not appear to have caused an increase in primate research, for instance.) Moreover, it is worth recalling that the 3Rs are ultimately ethical principles that, when first developed (Russell and Burch 1959), were thought to apply to invertebrates—and, in fact, are still thought to apply to invertebrates (National Research Council 2011). So, it is not clear that, even if ethics review for insects increased the use of vertebrate animal models in research, that would contradict 3Rs efforts. Indeed, it is not obvious how to think about the ethics of replacing smaller numbers of vertebrate animals with larger numbers of invertebrate animals, as often happens (Fischer 2025).

Ultimately, the broader commentary provided by respondents indicates a research community in transition. While some expressed caution about the idea of “insect welfare” and skepticism about ethics review, many also called for more research on insect welfare and more systematic ethics review, for the sake of public trust, scientific rigor, and institutional consistency. Overall, the evidence from this survey suggests that the UK entomological community is growing increasingly appreciative of ethics review and that ongoing development of evidence-based resources, communication, and training will be key to facilitating this progress as well as minimizing or avoiding the process and outcome concerns shared by survey participants.

## Supporting information

Supplemental File 1: University and institute list

Supplemental File 2: Paper version of the Question bank for insect welfare and ethics questionnaire

Supplemental File 3: Codebook

Supplemental File 4: RES membership

Supplemental File 5: Anonymised Responses

## Additional files

Supplemental File 1: University and institute list

Supplemental File 2: Paper version of the Question bank for insect welfare and ethics questionnaire

Supplemental File 3: Codebook

Supplemental File 4: RES membership

Supplemental File 5: Anonymised Responses

## Acknowledgements

The authors would like to thank the scientists that took time to complete the survey. We would also like to thank the Royal Entomological Society team that supported the project and advertised the survey. We would like to thank the Human Behaviour Change for Life for undertaking ethical review of the survey prior to launch and the three anonymous reviewers who provided valuable feedback.

This project was funded by a grant provided by Open Philanthropy to the IWRS; the funder did not have any influence over survey design, data analysis, manuscript preparation, or decision to publish the results. Any opinions, findings, and conclusions or recommendations expressed in this material are those of the authors and do not necessarily reflect the views of Open Philanthropy.

Resources from the project are freely available on https://www.royensoc.co.uk/insect-welfare-and-best-practices/

## Authors’ contributions

JES led the study design, analysis, wrote, edited and approved submissions. MB and BF supervised, wrote, edited and approved the manuscript; MB also analysed data. All authors read and approved the final manuscript.

## References

Aberystwyth University. (2025). Ethical review procedure for use of animals in research. Research, Business & Innovation. https://www.aber.ac.uk/en/rbi/support-services/ethics/animals/

Andrews, K., Birch, J., Sebo, J., and Sims, T. (2024). New York Declaration on Animal Consciousness. nydeclaration.com.

ASAB Ethical Committee/ABS Animal Care Committee. (2024). Guidelines for the ethical treatment of nonhuman animals in behavioural research and teaching. Animal Behaviour, 207, I–XI. 10.1016/S0003-3472(23)00317-2

Barrett, M., & Fischer, B. (2023). Challenges in farmed insect welfare: Beyond the question of sentience. Animal Welfare, 32, e4.

Barrett, M., Drewery, M., & Fischer, B. (2025). Entomologists’ knowledge of, and attitudes towards, insect welfare in research and education. Ecological Entomology, 50(1), 45–62.

Birch, J. (2017). Animal sentience and the precautionary principle. Animal Sentience, 16(1), 1–15.

Birch, J., Burn, C., Schnell, A., Browning, H., & Crump, A. (2021). Review of the evidence of sentience in cephalopod molluscs and decapod crustaceans. London School of Economics and Political Science.

Budda ML, Pritt SL. Evaluating IACUCs: Previous Research and Future Directions. J Am Assoc Lab Anim Sci. 2020;59(6):656–64.

Crump, A., Gibbons, M., Barrett, M., Birch, J., & Chittka, L. (2023). Is it time for insect researchers to consider their subjects’ welfare? PLOS Biology, 21(5), e3002138.

Crozier, G. K. D., & Schulte-Hostedde, A. I. (2015). Towards improving the ethics of ecological research. Science and Engineering Ethics, 21(3), 577–594.

Drinkwater, E., Robinson, E. J., & Hart, A.G. (2019). Keeping invertebrate research ethical in a landscape of shifting public opinion. Methods in Ecology and Evolution, 10(8), 1265–1273.

Field, K. A., Paquet, P. C., Artelle, K., Proulx, G., Brook, R. K., & Darimont, C.T. (2019). Publication reform to safeguard wildlife from researcher harm. PLoS Biology, 17(4), e3000193.

Fischer, B. (2025). Should Researchers Use Fruit Flies Instead of Mice in Pain Research?. Animal Behaviour and Welfare Cases, (2025), abwcases20250012.

Freelance, C.B. (2019). To Regulate or Not to Regulate? The Future of Animal Ethics in Experimental Research with Insects. Science and Engineering Ethics, 25 (5):1339–1355.

Gibbons, M., Crump, A., Barrett, M., Sarlak, S., Birch, J., & Chittka, L. (2022). Can insects feel pain? A review of the neural and behavioural evidence. Advances in Insect Physiology, 63, 155–229.

Harvey-Clark, C. (2011). IACUC challenges in invertebrate research. Ilar Journal, 52(2), 213–220.

IWRS. (2023.) About Insect Welfare. https://www.insectwelfare.com/insect-welfare

Jörgensen S, Lindsjö J, Weber EM, Röcklinsberg H. Reviewing the Review: A Pilot Study of the Ethical Review Process of Animal Research in Sweden. Animals. 2021; 11(3):708.

Loss, C. M., Melleu, F. F., Domingues, K., Lino-de-Oliveira, C., & Viola, G.G. (2021). Combining animal welfare with experimental rigor to improve reproducibility in behavioral neuroscience. Frontiers in Behavioral Neuroscience, 15, 763428.

Low, P. (2012). The Cambridge Declaration on Consciousness. Proceedings of the Francis CrickMemorial Conference, Churchill College, Cambridge University, July 7 2012, pp 1-2.

Milford A, De Clercq E, Louis-Maerten E, Geneviève LD, Elger BS (2025) How animal ethics committees make decisions – a scoping review of empirical studies. PLoS ONE 20(3):e0318570.

National Research Council. (2011). Guide for the care and use of laboratory animals (8th ed.). The National Academies Press. 10.17226/12910

Page, S. A., & Nyeboer, J. (2017). Improving the process of research ethics review. Research Integrity and Peer Review, 2(1), 14. 10.1186/s41073-017-0038-7

Perl CD, Kissinger C, Godfrey RK, Castillo EM, Fischer B, Barrett M (2025) Identifying trends in reporting on the ethical treatment of insects in research. PLoS One 20(8):e0328931. 10.1371/journal.pone.0328931

Pritt, S. L., & Hammer, R. E. (2017). The interplay of ethics, animal welfare, and IACUC oversight on the reproducibility of animal studies. Comparative Medicine, 67(2), 101–105.

R Core Team (2025), R: A language and environment for statistical computing. R Foundation for Statistical Computing, Vienna, Austria. https://www.R-project.org/.

Rethink Priorities. (2021). Submission of evidence to Animal Welfare (Sentience) Bill. UK Parliament. https://committees.parliament.uk/writtenevidence/37632/html/

Royal Entomological Society. (2024, October 16). RES statement on the ethical treatment of insects. https://www.royensoc.co.uk/news/res-statement-on-the-ethical-treatment-of-insects/

Russell, W. M. S., & Burch, R. L. (1959). The principles of humane experimental technique. Methuen.

Schuppli, C. A. (2011). Decisions about the Use of Animals in Research: Ethical Reflection by Animal Ethics Committee Members. Anthrozoös, 24(4), 409–425.

Silverman J, Baker SP, Lidz CW. A self-assessment survey of the institutional animal care and use committee, part 1: animal welfare and protocol compliance (vol 41, pg 230, 2012). Lab Anim. 2012;41(8):230–5. pmid:22821046

Trout, R. T., Minteer, C. R., Pallipparambil, G. R., Magnus, R. M., & Wiedenmann, R.N. (2010). A perspective on education in research ethics for entomology graduate students. American Entomologist, 56(4), 198.

Varga O. Critical Analysis of Assessment Studies of the Animal Ethics Review Process. Animals (Basel). 2013;3(3):907–22.

