## Supplemental File 2: Paper version of the Question bank for insect welfare and ethics questionnaire for "Ethics review for insect research in the UK: Patterns, drivers, and perceived impact on research quality and process"

### Background statement

Thank you for participating in this survey. It should take you approximately 15-35 minutes to complete. We are aiming to gather insights from scientists that work with insects regarding their experiences of the ethical use of insects in research. Your input is vital for identifying knowledge gaps and improving the support available for researchers in this field.

This questionnaire will cover various aspects of the ethics application process, including submission practices, perceived challenges, and attitudes toward ethical oversight in entomological research. The findings will contribute to a better understanding of the needs and experiences of researchers and may help inform future resources and policies.

*Ethics statement*

This study has undergone ethical review by the Human Behaviour Change for Life (HBCL) Organisation. This survey is produced by the Royal Entomological Society, using Survey Monkey software. We are following the same online survey privacy statement as the HBCL which can be found here: https://hbcforlife.org/online-survey-privacy-statement/ .

*Confidentiality statement*

We value your privacy and are committed to protecting your personal information. All responses to this survey will be kept confidential and will be used solely for research purposes. Individual responses will not be identifiable in any reports or publications. Data will be aggregated and anonymised to ensure that no individual participant can be recognised.

*Right to remove data.*

Your participation in this survey is entirely voluntary. You have the right to withdraw from the survey at any time without any negative consequences. Additionally, you can request the removal of your data from the study at any point prior to the questionnaire closing date by contacting Dr Jessica

Stokes at the Royal Entomological Society (**). Upon your request, we will ensure that your data is deleted from our records. After the questionnaire closing date all data will be anonymised.

*Participation Statement*

I have read the above and understand what the research involves.

I understand my participation in the research is voluntary, and if I decide that I no longer wish to take part in this research, I should not submit my response.

I am over the age of 18.

I consent to the processing of my personal information for the purposes of this research.

I understand that such information will be treated as strictly confidential and handled in accordance with the provisions of the Data Protection Act 2018.

I agree that the project has been explained to me to my satisfaction and I agree to take part in this research.

I am not suffering from any health conditions that would prevent me from taking part in this research.

I understand that the information I have submitted will aid in the objectives of the project as outlined in the project introduction or explanation already giving to me. Confidentiality and anonymity will be maintained and it will not be possible to identify me from any publications, unless I have given expressed permission.

Thank you for your valuable contribution to this important research.

Royal Entomological Society

### Background questions

1. Title
2. Name
3. Institution
4. Is your research currently based in the UK?
    a. Yes
    b. No

    *(If No skip to debrief)*
5. What is your current position?

    - Undergraduate Student
    - Masters student
    - PhD student
    - Postdoc/ ECR (less than 5 years post PhD)
    - Senior Postdoc/Fellow/Reader
    - Professor
    - Emeritus Professor/ Retired
    - Research Assistant/ Technician
    - Industry
    - Zoo or insectarium
    - Other, please describe

6. How many years have you been involved in entomological research?

    - <1
    - 1-2
    - 3-5
    - 6-10
    - 11-20
    - 20+

Royal Entomological Society

7. What is your primary area of research?

    - Agricultural and forest entomology
    - Behaviour and cognition
    - Biomedical (non-insect genomics, pharmacology)
    - Biotechnology (insect derived products, waste management)
    - Conservation and diversity
    - Ecological
    - Education and outreach
    - Forensic entomology
    - Insect neuroscience and physiology
    - Molecular biology (inclusive of insect genomics)
    - Vector and pathogen (medical, veterinary, urban)
    - Other, please describe

8. What is your primary research species?
   (Open question)

*(All answers proceed to Question 9)*

### Current Ethics application status

9. Do you currently submit ethical applications for your work with insects to your internal ethics review committee (AWERB)?

    a. No, I do not current submit ethics applications for my work with insects *(skip to Qu 20 Attitudes section)*
    b. Yes, I am required to submit ethics applications for my work with insects (it is mandatory) *(Continue to Qu 10)*
    c. Yes, I voluntarily submit ethics applications for my work with insects (it is not mandatory) *(Continue to Qu 10)*

*(Answers B & C from Question 9 proceed to Question 10 (Branches B and C respectively), Answers A from Question 9 proceed to Question 21 (Branch A))*

10. Roughly how many ethics applications have you submitted in the past year? (include those of your students where you have had supervisory oversight or helped draft the application)

    - 0
    - 1-2
    - 3-5
    - 6-10
    - 11-20
    - 20+

*(Only Branch B answers proceed to Question 11)*

11. When did you start submitting mandatory ethics application for your research involving insects?

- This year (2024)
- 2023
- 2022
- 2021
- 2020
- (continue dropdown list to 1986)

*(Branch B & C proceed to question 12)*

### Experiences

12. How confident do you feel about the ethics application process?
    a. Not confident at all
    b. Somewhat not confident
    c. Neutral
    d. Somewhat confident
    e. Very confident

13. How confident are you that you can deliver what is expected of you when completing an ethics application?
    a. Not confident at all
    b. Somewhat not confident
    c. Neutral
    d. Somewhat confident
    e. Very confident

14. What feedback have you received from your internal ethics review committee (AWERB) regarding your application? (Please check all that apply)
    a. Positive comments
    b. Suggestions for improvement of study methodology
    c. Rejections and reasons provided
    d. No feedback received
    e. Other (please specify):

*15.* Can you provide any examples of the feedback you have received from your ethics applications (Optional): (Open comment box)

### Resources & Support

16. What resources or support do you think would help you (and/or your students) with ethics applications submissions? (Please check all that apply)
    a. Undergraduate curricular materials on insect welfare
    b. Training on the ethical use of animals in research with examples of how to apply this to insects
    c. Guidance on how to develop an ethical application for submission to ethics panels (AWERBs)
    d. Greater research on insect welfare
    e. Information on insect biology for those on the ethics panels (AWERB members)
    f. Other, please comment

17. Have you utilised any external resources or tools that have been particularly helpful in preparing submissions for ethical applications? If so, what were they? (Optional)

*(Branches B and C proceed to Attitudes questions)*

### Attitudes

18. Which of the following factors influence your decision to submit an ethics application for your insect research? (Please select all that apply)
    a. Institutional requirements
    b. Personal ethical beliefs
    c. To ensure the welfare of research subjects
    d. Application of best practice methodologies and standardisation of research
    e. To receive feedback on study design from ethics panel
    f. Grant requirements
    g. Requirement to publish
    h. To address concerns about public perception of my research practices
    i. Comments (any other reasons, please list):

19. How valuable do you consider ethics applications to be in your entomological research?
    a. Very important
    b. Important
    c. Neutral
    d. Unimportant
    e. Very unimportant
    f. Comments

20. Have any of your practices changed as a result of the ethics application process?
    a. Yes (please comment, optional)
    b. No
    c. Unsure
    d. Comments

*(Branch A rejoins)*

21. In your opinion, how does the ethics application process influence the quality of research?
    a. It improves quality
    b. It has no effect
    c. It detracts from quality
    d. Comments

### Feedback

22. Please share any additional comments or insights regarding your experience with, or attitudes about, the ethics application process in entomological research

Royal Entomological Society

### Debrief

Thank you for participating in our survey regarding ethics applications in entomological research. Your input is invaluable and will contribute significantly to our understanding of the current landscape of ethics review processes within the field.

*What Happens Next*

Responses will be collated for anonymity and collectively analysed to identify trends and insights. The findings will be shared by the Royal Entomological Society in academic publications and presentations, helping to inform policy and practice in research ethics. We assure you that all data will be treated confidentially, and individual responses will remain anonymous.

Please remember that participation is entirely voluntary, and you have the right to withdraw until the closing date of the questionnaire, at which point the data will be anonymised and we will be unable to identify your data. If you have any questions or would like to withdraw your data, please contact our lead researcher Dr Jessica Stokes at.

*Additional Information*

If you have any further questions about the study, its objectives, or your rights as a participant, please feel free to reach out. We appreciate your time and effort in contributing to this important research.

Thank you once again for your valuable contribution!

*Resources*

https://www.insectwelfare.com/power-analysis-guide
https://www.insectwelfare.com/research-guidelines
https://www.royensoc.co.uk/news/res-statement-on-the-ethical-treatment-of-insects/
https://www.asab.org/ethics

Royal
Entomological
Society
