## Supplemental File 4: RES membership for "Ethics review for insect research in the UK: Patterns, drivers, and perceived impact on research quality and process"

| Membership Type | Total | Proportion of Total Membership (%) |
|---|---|---|
| Student | 573 | 24.5 |
| Regular | 502 | 21.4 |
| Retired | 365 | 15.6 |
| **Total RES Membership** | **2344** | |

**Supplemental File 4 Table 1. Royal Entomological Society (RES) Membership Demographics by Career Stage (2025).** Demographic data provided by RES in 2025. Other membership categories not shown that make up the total membership include staff membership, fellows, overseas and honorary members.
